# ViReaDB: A user-friendly database for compactly storing viral sequence data and rapidly computing consensus genome sequences

**DOI:** 10.1101/2022.10.21.513318

**Authors:** Niema Moshiri

## Abstract

**Motivation:** In viral molecular epidemiology, reconstruction of consensus genomes from sequence data is critical for tracking mutations and variants of concern. However, storage of the raw sequence data can become prohibitively large, and computing consensus genome from sequence data can be slow and requires bioinformatics expertise.

**Results:** ViReaDB is a user-friendly database system for compactly storing viral sequence data and rapidly computing consensus genome sequences. From a dataset of 1 million trimmed mapped SARS-CoV-2 reads, it is able to compute the base counts and the consensus genome in 16 minutes, store the reads alongside the base counts and consensus in 50 MB, and optionally store just the base counts and consensus (without the reads) in 300 KB.

**Availability:** ViReaDB is freely available on PyPI (https://pypi.org/project/vireadb) and on GitHub (https://github.com/niemasd/ViReaDB) as an open-source Python software project.

## 1 Introduction

Viral molecular surveillance, a technique in which viral genomes are reconstructed from sequence data generated by samples collected from patients as well as the environment (e.g. wastewater) and are monitored in real-time or near real-time, has been critical throughout the COVID-19 pandemic (Oude Munnink *et al.*, 2021; Karthikeyan *et al.*, 2022). The reconstruction of consensus genome sequences from raw sequence data requires the use of various bioinformatics pipelines, which can be slow and can require non-trivial computational expertise (Truong Nguyen *et al.*, 2021; Posada-Céspedes *et al.*, 2021; Moshiri *et al.*, 2022).

To minimize the need for redundant reanalysis of raw data, final consensus genomes can be stored in databases such as GISAID, which currently stores over 13.5 million SARS-CoV-2 genome sequences, over 5.5 million of which are complete genomes with high coverage (Shu & McCauley, 2017). However, the GISAID database, which is currently the most prominent database to which researchers submit SARS-CoV-2 and Monkeypox genome sequences, relies on the submitter to verify the validity of the genome sequence. Because the database does not store raw sequence data or base counts (likely due to storage limitations), errors in genome sequences resulting from bioinformatics errors cannot be detected and corrected purely from data stored within the database. Instead, individuals must request raw data directly from the submitting lab (Pekar *et al.*, 2022).

Here, we introduce ViReaDB, a user-friendly database for compactly storing viral sequence data and rapidly computing consensus genome sequences. By providing a user-friendly interface to rapidly compute base counts and consensus genomes directly within the database itself, ViReaDB aims to minimize the potential for bioinformatics error, and by storing raw data and base counts alongside the final consensus genomes, ViReaDB aims to improve reproducibility and validation of viral consensus genome sequences.

## 2 Results and discussion

ViReaDB is a cross-platform database interface written in Python 3 and depends on Pysam (Bonfield *et al.*, 2021) and NumPy (Harris *et al.*, 2020). ViReaDB optionally depends on Samtools (Li *et al.*, 2009) to add support for input files in the SAM or BAM format, and it optionally depends on Minimap2 (Li, 2018) to add support for input files in the FASTQ format (which ViReaDB will map to the reference genome).

The underlying ViReaDB database format is a SQLite database with two tables: “meta” and “seqs”. The “meta” table stores metadata about the database: the ViReaDB version number that created it as well as the reference genome name, sequence, and Minimap2 index (if Minimap2 is installed). The “seqs” table stores the viral sequence data added to the database, where each entry has the following items: the raw sequence data stored as an LZMA-compressed CRAM 3.0 file (Fritz *et al.*, 2011), an LZMA-compressed NumPy array containing base counts at all positions of the reference genome, an LZMA-compressed JSON containing insertion counts, and the LZMA-compressed consensus genome sequenced.

A new ViReaDB can be created via **create_db**(`db_fn,ref_fn`), where db_fn is the desired output SQLite file, and `ref_fn` is the reference genome FASTA. Alternatively, an existing ViReaDB can be loaded via **load_db**(`db_fn`), or multiple ViReaDBs can be merged via **merge_dbs**(`out_db_fn, in_db_fns`), where in_db_fns is a list of input ViReaDB SQLite files. The user can then add a new entry to a ViReaDB via **add_entry**(`ID, reads_fn`), where ID is the unique identifier of this sample, and reads_fn is the raw sequence data of this sample in the CRAM, BAM, SAM, or FASTQ format. CRAM files are added as-is, BAM and SAM files are converted to LZMA-compressed CRAM using Samtools, and FASTQ files are mapped to the reference genome using Minimap2 and are converted to LZMA-compressed CRAM using Samtools. After an entry is added to a ViReaDB, base counts and insertion counts can be computed from the CRAM using Pysam via **compute_counts**(`ID`): counters are pre-allocated for all positions of the reference genome, and read mappings are processed on-the-fly to increment the corresponding base counts of the positions covered by the read and to increment insertion counts. Once base and insertion counts have been computed, which is the most computationally-expensive aspect of ViReaDB, a consensus genome can be computed from the counts via **compute_consensus**(`ID`). Comprehensive usage details can be found in the ViReaDB documentation, and example commands can be found in the ViReaDB Cookbook, both of which can be found on the GitHub repository.

Because of LZMA compression, base counts, insertion counts, and consensus genome sequences are extremely small when stored in a ViReaDB and are constant in file size with respect to sequencing depth (they scale linearly with reference genome length): the vast majority of ViReaDB file size is due to the raw reads. In order to provide a highly-compact manner in which data can be shared, ViReaDB allows the user to remove reads from a given entry, resulting in just base/insertion counts and consensus genome sequence that occupy less than 400 KB total. This provides a middle-ground for data sharing: rather than being forced to choose between sharing final consensus genome sequences (which provide no contextual information about the raw data) or sharing the raw CRAM/BAM/SAM/FASTQ data (which can be prohibitively large), users can share the consensus sequence packaged with base/insertion counts to provide context about coverage and confidence in base calls, but without the burden of large file sizes.

In order to benchmark ViReaDB with respect to file size runtime, we obtained a SARS-CoV-2 amplicon sequencing dataset in which 2,607 samples were sequenced PE150 across four lanes of an S4 flow cell to an average read count of 4.58 M read pairs per sample using the SWIFT v2 protocol on an Illumina NovaSeq 6000 (Moshiri *et al.*, 2022). Samples were mapped to the NC_045512.2 reference genome using Minimap2 and were trimmed using iVar Trim (Grubaugh *et al.*, 2019). We selected the single highest-depth sample and randomly subsampled it to *n* = 100, 1K, 10K, 100K, and 1M successfully-mapped reads, with 10 replicates for each *n*. We then built ViReaDB databases, computed base/insertion counts, and computed consensus sequences from each subsampled replicate.

When including reads, ViReaDB file size grows roughly linearly for large *n*, and a ViReaDB storing 1 million mapped reads is ~50 MB. After computing base/insertion counts and consensus genomes, removing reads from the ViReaDB results in files smaller than 400 KB. Computing base/insertion counts scales roughly linearly with the number of mapped reads and takes ~660 seconds (~11 minutes) on a dataset of 1 million reads. Computing the consensus genome from base/insertion counts takes less than 1 second, even with 1 million reads (Fig. 1).

**Fig. 1.**
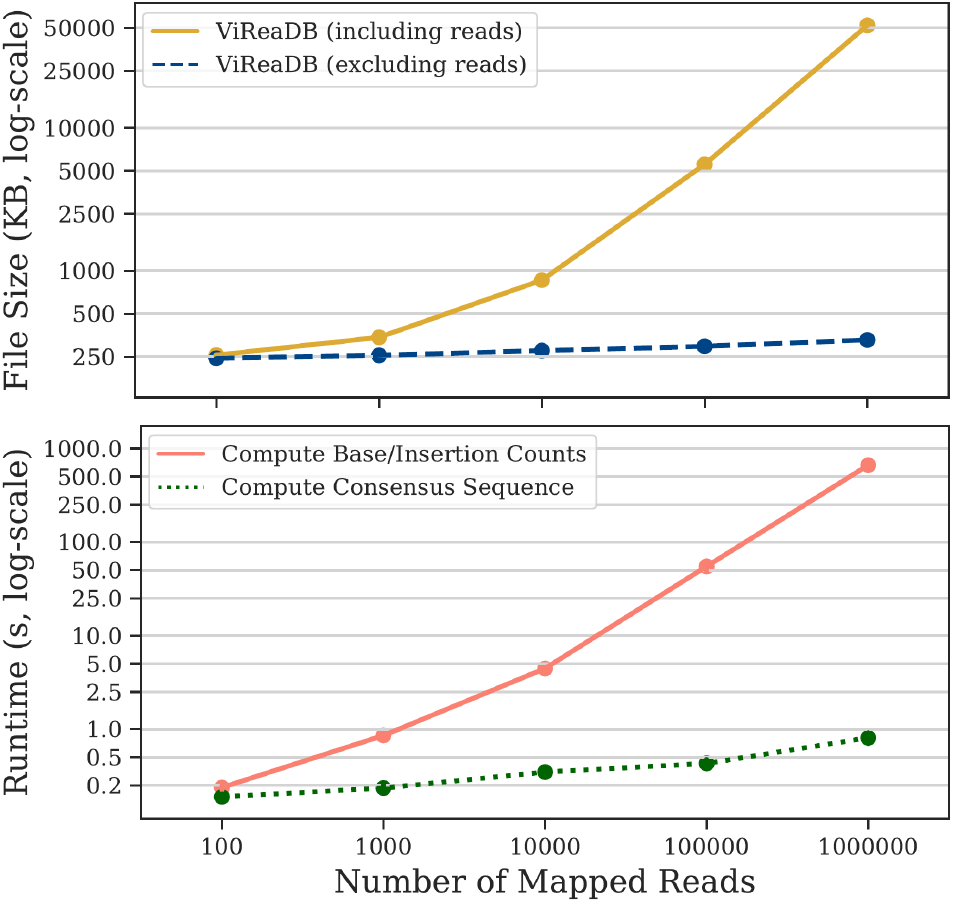
Benchmark results. File size (top) and execution time (bottom) for SARS-CoV-2 sequence datasets with *n* = 100, 1K, 10K, 100K, and 1M mapped reads. All runs were executed sequentially on a 2.8 GHz Intel i7-1165G7 CPU with 16 GB of memory.

Future ViReaDB updates will (1) migrate to CRAM 3.1 and 4.0 upon official release for improved compression (Bonfield, 2022), and (2) implement multithreading in computing base/insertion counts.

## Acknowledgements

We would like to thank Rob Knight, Louise Laurent, Kristian Andersen, and Karthik Gangavarapu for fruitful conversations.

## Funding

This work has been supported by UC San Diego faculty research funds.

## Conflict of Interest

none declared.

## Notes

https://github.com/niemasd/ViReaDB

https://github.com/niemasd/ViReaDB-Paper

### Competing Interest Statement

The authors have declared no competing interest.

